# *resLens*: genomic language models to enhance antibiotic resistance gene detection

**DOI:** 10.1101/2025.07.08.663767

**Authors:** Matthew Mollerus, Katharina Dittmar, Keith A. Crandall, Ali Rahnavard

## Abstract

The rise of antibiotic resistance necessitates advanced tools to detect and analyze antibiotic resistance genes (ARGs). We present *resLens*, a family of genomic language models that leverage latent genomic representations to enhance ARG detection and analysis. Unlike alignment-based methods constrained by reference databases, *resLens* fine-tunes a pre-trained DNA language model on curated ARG datasets, achieving competitive or superior performance in classifying resistance genes across multiple evaluation scenarios, including when ARGs exhibit sequences and mechanisms of resistance dissimilar to those in reference datasets.

## Introduction

Rising levels of antibiotic resistance in microbial pathogens require new tools to study the presence and evolution of antibiotic resistance genes (ARGs). We propose *resLens*, a novel genomic language model that harnesses its latent understanding of genomic data to improve detection of ARGs from metagenomic data and identify potential novel ARGs.

Most existing tools are alignment-based, either as best hit algorithms such as ResFinder,^1^ ARG-ANNOT,^2^ AMR++,^3^ and RGI,^4^ k-mer approaches such as KARGA,^5^ Hidden Markov Model algorithms such as Meta-MARC,^6^ or using alignment-based features for machine learning models, such as DeepARG.^7^ These approaches are limited in a variety of ways. First, they perform poorly when variants do not closely match reference ARGs in their database. Second, the databases represent a small fraction of the resistome and can struggle to keep up with the rapid pace of resistance evolution.^8^ Finally, reference database approaches are incapable of identifying substantially novel genes or mutations that confer resistance.^9,10^ Other efforts to predict a resistance phenotype from an underlying genotype rely on some combination of whole genome data,^11,12^ species-specific models,^13^ or causal information about the resistance evolution to predict novel genes,^14^ all of which make them have limited applicability to many real-world applications.

Several deep learning models have attempted to overcome these limitations by using neural networks to learn functional representations of ARGs. ARGNet trained autoencoders on one-hot representations of amino acid sequences of ARGs to embed them in a latent space then trained convolutional neural networks to predict ARG class from those embeddings.^15^ HMD-ARG similarly inputs one-hot representations of amino acid sequences into a set of convolutional neural networks.^16^ These methods are more dynamic than alignment-based algorithms but must build their representations of ARGs and protein function from scratch.

Transfer learning approaches borrowed from the field of natural language processing offer to bridge this gap. Applying large language model architectures, specifically transformer encoder models,^17^ to DNA sequence data has yielded models with a latent understanding of genomic elements and state-of-the-art performance of a variety of genomic classification tasks.^18,19^ These models divide a DNA sequence into small subsequences called tokens, embed them in a high-dimensional space, and then update the token embeddings based on the context of surrounding sequence to generate a final output embedding that can be used for unsupervised or supervised analysis. These embeddings encode information about both sequence identity and biological function.^20^ The models are first pre-trained on large datasets of whole genomes via masked language modeling to develop a general understanding of the relationship between genomic elements and then are optionally fine-tuned on smaller, task-specific labeled datasets (**Fig. 1b**).

**Figure 1.**
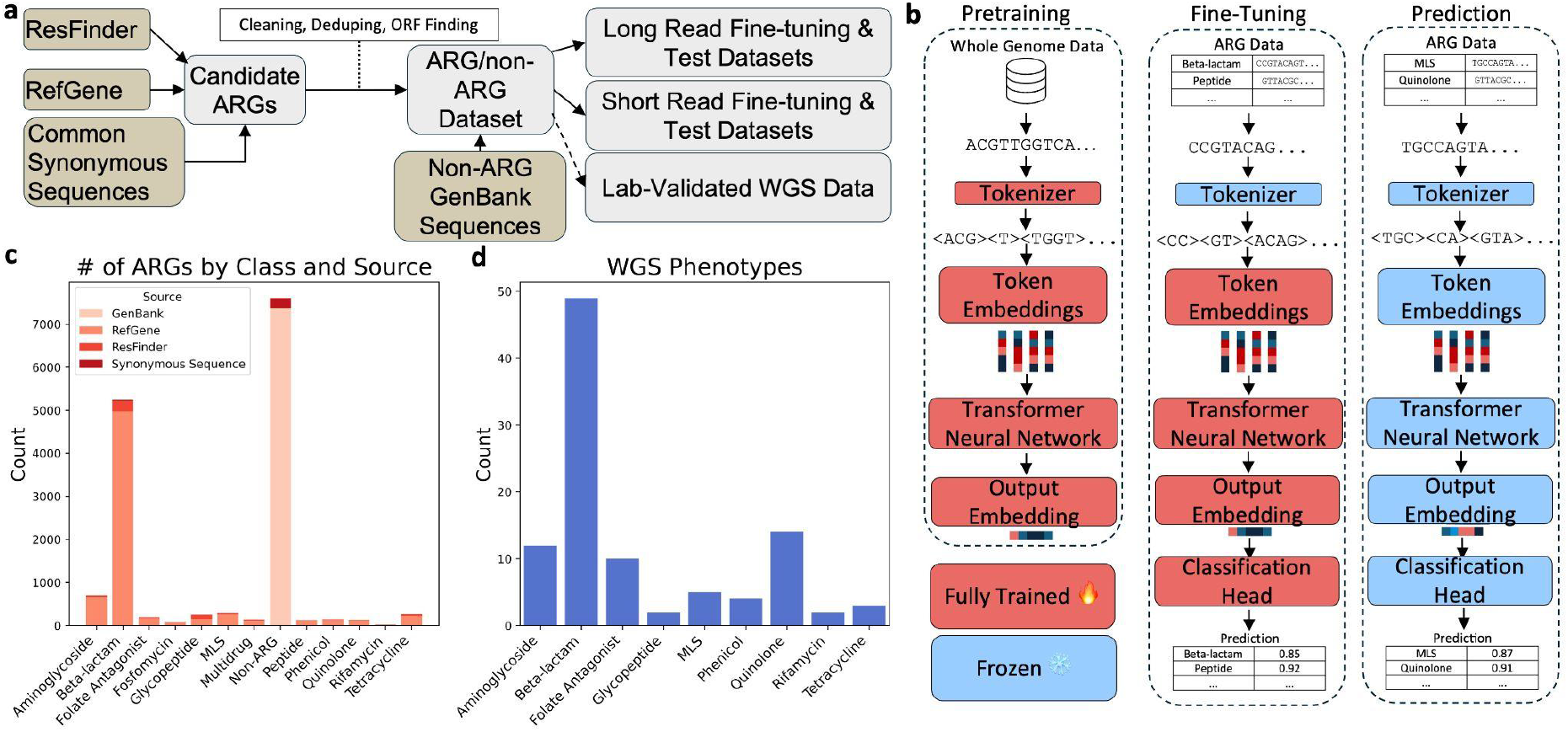
Data and modelling overview. **a**, Flow chart of construction and application of datasets. **b**, Overview of gLM pretraining, fine-tuning, and inference, with frozen and trainable layers at each step displayed. **c**, Distribution of phenotypes presented in LR training and test data, colored by source. **d**, Distribution of phenotypes presented in lab-validated Whole Genome Sequencing (WGS) data.

*resLens* fine-tunes a seqLens 89M parameter pre-trained DNA language model on a broad dataset of resistance genes, using the output embeddings to classify input sequences as conferring or not conferring resistance to different classes of antibiotics.^21^ We constructed models to classify long read (LR) and short read (SR) data and evaluated it and several other tools used to identify ARGs against held out test sets. Additionally, we performed two sets of tests to evaluate resLens’s ability to identify ARGs that are novel or substantially dissimilar to its training data. First, we retrained the model with two subclasses of ARGs held out and then evaluated the models’ ability to classify them correctly; second, we remade the train and test splits by first clustering by sequence similarity then assigning different clusters to splits, allowing us to quantify the models’ performance on out-of-sample data. Finally, we demonstrated a potential usage of *resLens* to identify genes conferring resistance in assembled genomes of organisms with lab-validate resistance phenotypes.

## Results

### Train and Test Datasets

The fine-tuning datasets consisted of known ARGs from ResFinder and the NCBI Pathogen Detection RefGene database,^22^ as well as synonymous nucleotide variants for some of the most common ARGs. We kept only one instance of perfect duplicates and the longer genes in the case that one gene was a perfect subsequence of another, then kept only genes that conferred resistance to antibiotic classes with at least 20 instances. We then passed the sequences through Prodigal^23^ to ensure they contained only open reading frames. This preprocessing yielded 7,606 ARGs over 12 classes of antibiotics. This is fewer classes than many tools classify, as merging databases required consolidating more specific classes into broader parent classes. We combined these ARGs with an equal number of non-resistance bacterial gene sequences of similar length, randomly selected from GenBank, filtered to ensure no more than 90% sequence identity with any ARG sequences, and processed with the same ORF finding process. We created an additional short-read dataset by splitting up the whole gene sequences into 150 bp reads. We finetuned four models in total, two each for both the LR and SR data. For each dataset, one model first performed a binary classification of ARG or non-ARG, then a second model classified the predicted ARGs into specific ARG classes. In each case, we split the dataset into 80% train and 20% test data, then performed 10-fold cross validation with the train dataset, and then evaluated the best model checkpoint from cross validation via weighted F1 on the test set. In cases where the sequence was longer than the model’s context window, it was split up into two or more shorter sequences, all of which inherited the same label (**Fig. 1a**).

On each test dataset, we evaluated the appropriate *resLens* models against two other deep learning models (ARGNet and DeepARG), and five broadly alignment-based tools (RGI, KARGA, Meta-MARC, ResFinder, and AMR++). We evaluated each tool using the default parameters for the type of data unless there was a compelling reason to change them; see the Methods section for details of the usage of each tool. All tools except AMR++ provided read-level labels and were evaluated with weighted F1, MCC, precision, and recall; to compare AMR++, we computed the Jensen-Shannon divergence and Spearman correlation of the distribution of labels in the test set and that of the predictions of AMR++, *resLens*, and selected other tools.

*resLens* performed better than other methods on the LR test dataset, although the difference between it, RGI including loose hits, RGI excluding loose hits, and KARGA were modest (weighted F1 scores of 0.9690, 0.9686, 0.9650, and 0.9602 respectively) (**Fig. 2a**). On the SR test dataset, RGI including loose hits and KARGA meaningfully outperformed the *resLens* models, (weighted F1 scores of 0.9577, 0.9656, and 0.9155 respectively) (**Fig. 2b**). The *resLens* LR models also more closely reproduced the distribution of classes in the test dataset than the other models compared, including AMR++ (**Fig. 3a**). The *resLens* models demonstrated competitive inference times on the test set. On the LR test set, only ARGNet was faster than *resLens* by wall clock time (19.19s and 23.82s, respectively), while on the SR test set, KARGA and DeepARG were faster (15.31 and 25.74, respectively, vs. 37.61 for *resLens*) (**Fig. 3b**).

**Figure 2.**
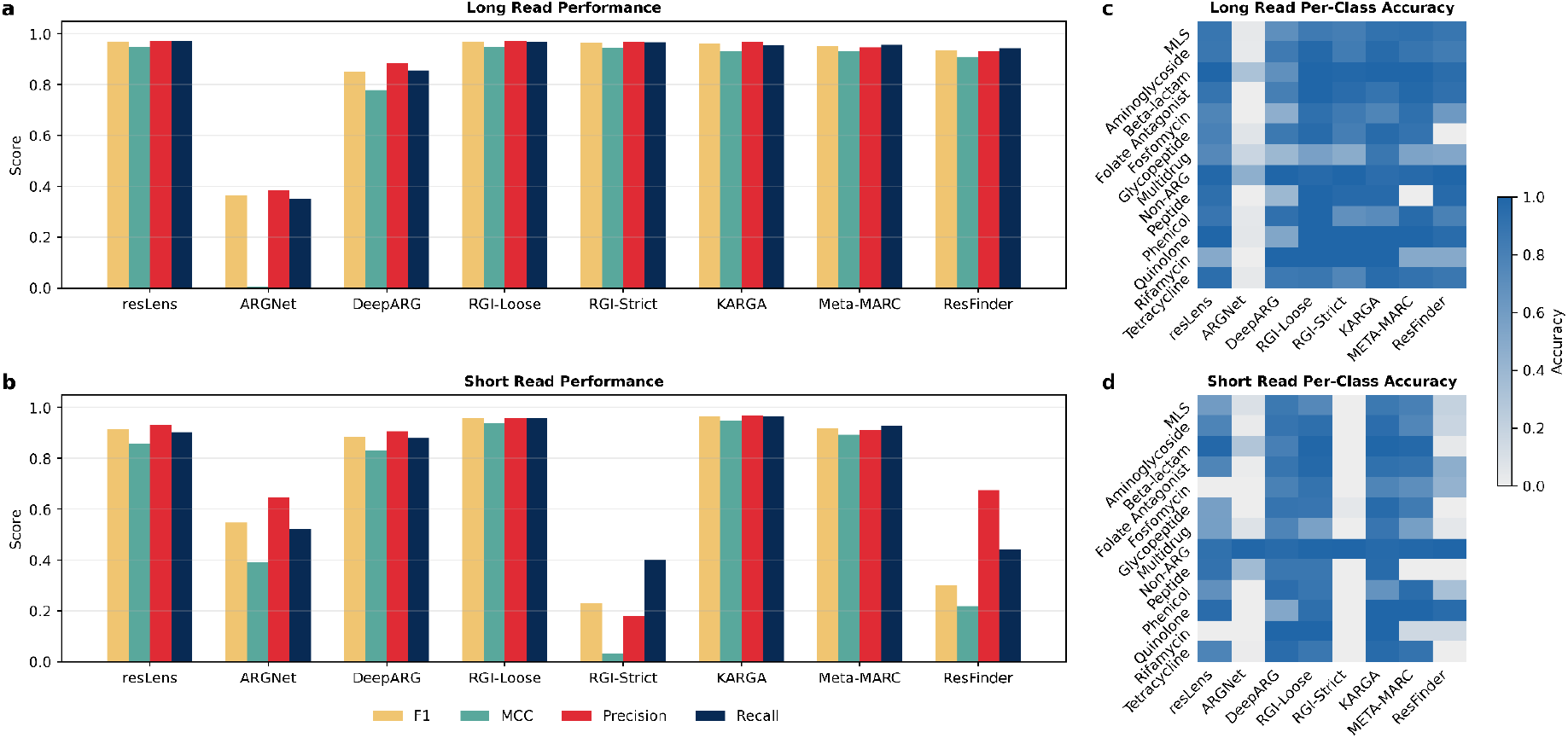
Comparison of classification performance across different tools and datasets. **a**, Bar plot showing weighted F1, MCC, precision, and recall scores for selected tools on the LR test set. **b**, Bar plot showing weighted F1, MCC, precision, and recall scores for selected tools on the SR test set. **c**, Heatmap of accuracy for each ARG class for each tool on the LR test set. **d**, Heatmap of accuracy for each ARG class for each tool on the SR test set.

**Figure 3.**
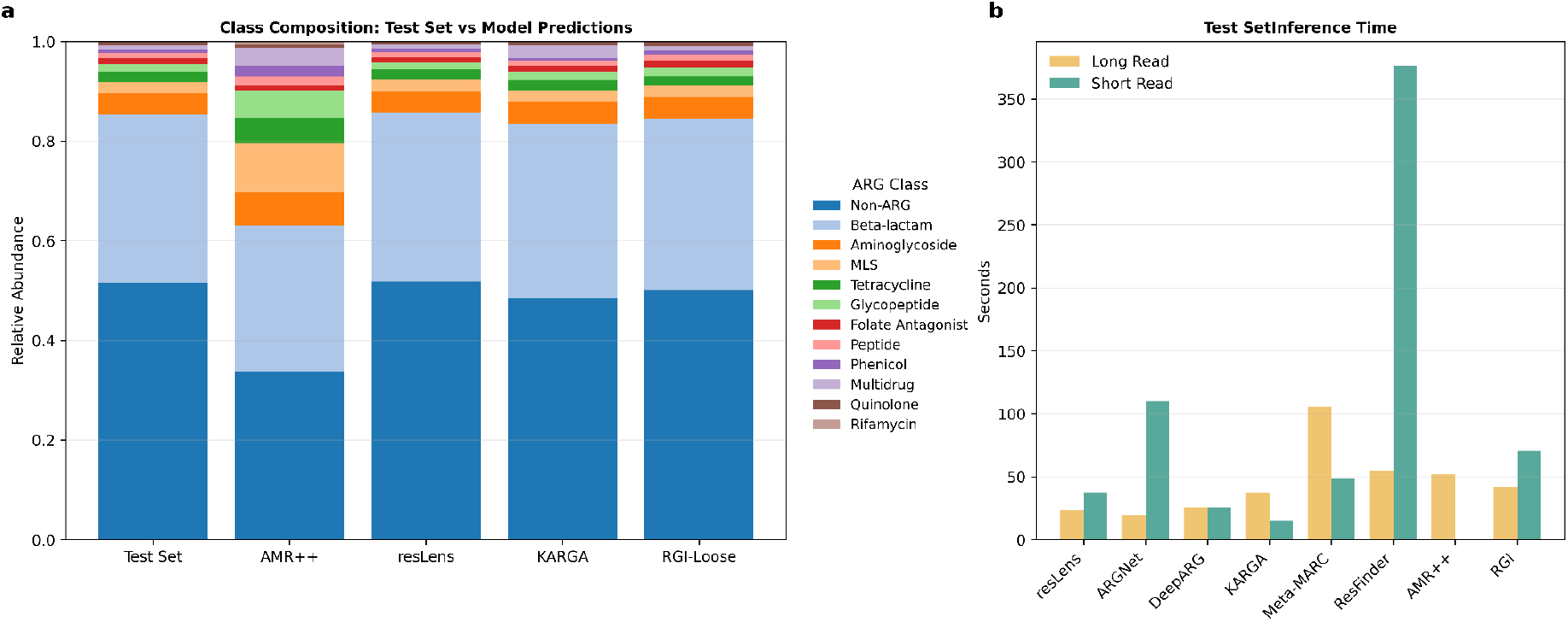
**a**, Stacked bar chart of composition of the LR test set against predicted composition by selected tools. This is useful for comparing tools that do not make gene-level predictions, such as AMR++. **b**, Bar plot of wall clock inference times on LR and SR test sets by tool.

### Novel ARG Classification

To evaluate the performance of the model on novel ARGs, we took advantage of the fact that many different families of genes can confer resistance to the same class of antibiotics. We identified two gene families, one that conferred resistance to beta-lactams and one that conferred resistance to aminoglycosides, that had low sequence similarity to other gene families that conferred resistance to the same antibiotics. For the beta-lactams, this group was the blaADC family of beta-lactamases, while for aminoglycosides this was the aminoglycoside nucleotidyltransferase (ANT) gene family; the blaADC sequences had a maximum of 61.6% identity with the closest other beta-lactam ARG sequence, while the ANT sequences had a maximum of 49.8% identity with the closest other aminoglycoside ARG sequence. We then created a new LR test dataset consisting of just those two gene families and a new LR train set consisting of all other genes, excluding those without gene family annotations to prevent data leakage. The pre-trained model was then fine-tuned with this new train dataset using ten-fold cross validation and evaluated on the new test set. It accurately classified the heldout genes, albeit at a lower rate for the ANT genes (100% for the blaADC genes in both cases, 84.7% vs 99.7% for ANT genes) than in the main stages of model development.

To fairly compare *resLens* to an alignment-based approach on this task, we created a version of the ResFinder database with the blaADC and ANT genes removed, then had ResFinder make predictions on the new test set of heldout gene families. It performed notably worse, identifying none of the blaADC genes and 86.03% of the ANT genes. Unfortunately, we were unable to fairly evaluate the other deep learning approaches on this task due to issues with reproduction.

We generalized this analysis by constructing another quartet of train and test datasets. We first use CD-HIT-EST to create clusters of genes that share no more than 90% identity with each other, then split the clustered into train and test splits, ensuring that no sequence in the test dataset was similar to any in the train dataset.^24^ We then fine-tuned new LR and SR models with the new train datasets and evaluated them on the new test datasets, using the same procedure as before.

This approach resulted in a decrease in performance when compared to the original models. The weighted F1 scores for the LR and SR model full predictions on the clustered test sets was 0.803 and 0.689, respectively. Almost all of the decrease in performance was concentrated in the binary models, as the weighted F1 scores for the LR binary and multiclass models dropped from 0.978 to 0.709 and from 0.977 to 0.944, respectively; the weighted F1 scores for the SR binary and multiclass models dropped from 0.873 to 0.704 and from 0.888 to 0.84,8 respectively. The per-class accuracies dropped in proportion to overall performance decrease (**Fig. 4a**). To visualize the difference in internal representations caused by the different splitting methods, we extracted the CLS tokens for the appropriate train and test datasets from the final layers of the LR multiclass model trained on random splits and from that trained on the clustered splits and constructed t-SNE plots of the token embeddings (**Fig. 4b**,**c**). In both plots, the train data points are tightly clustered by class with test data points generally falling within the train data of the same class; however, more test data points fall in clusters composed of data points of a different class for the model trained on clustered splits than random splits, which is indicative of increase in misclassification due to the distribution shift between train and test data in that context.

**Figure 4.**
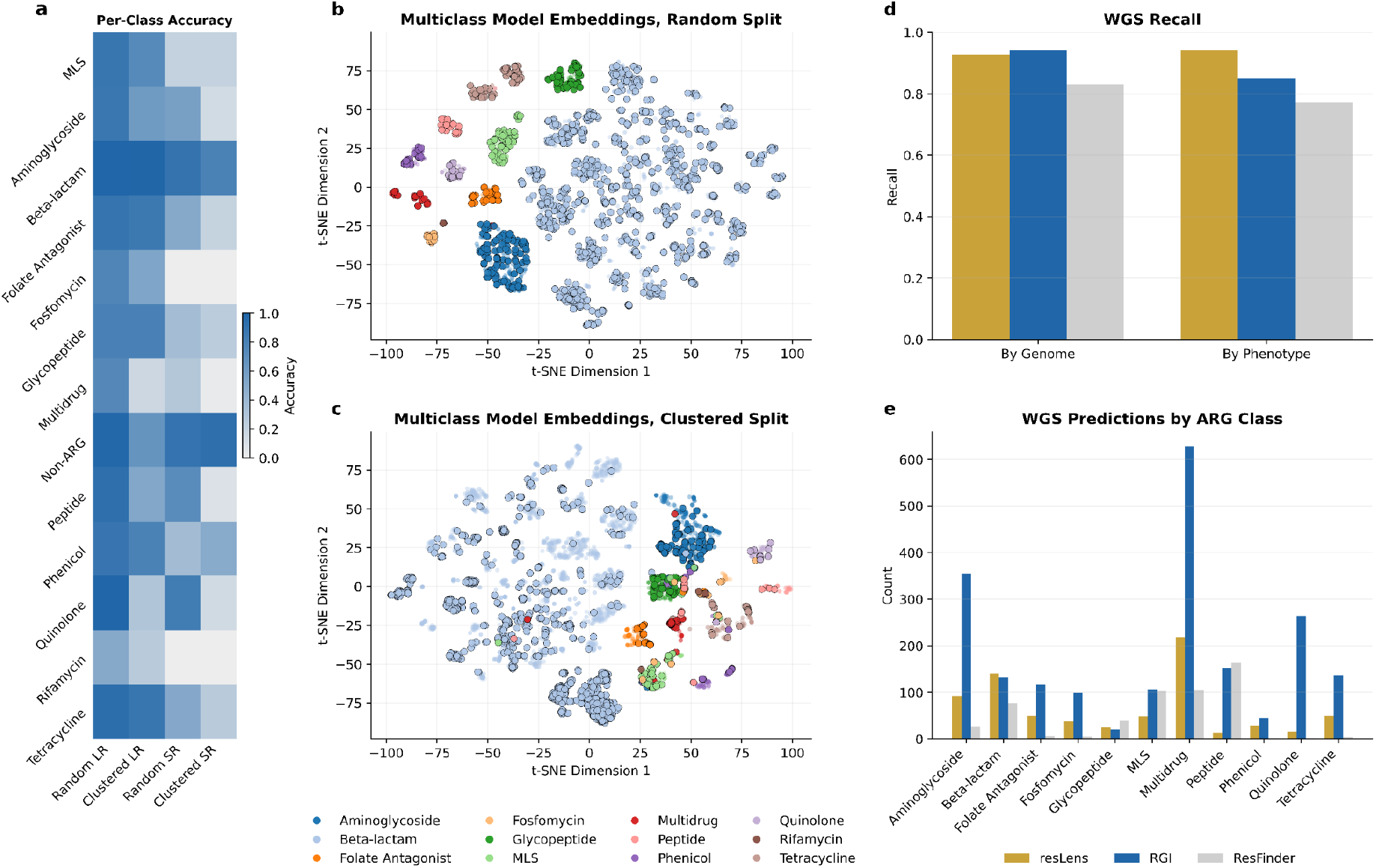
**a**, Heatmap of accuracy by class for *resLens* models trained with random splits and clustered splits. **b**, t-SNE plot of CLS token embeddings for LR Multiclass model trained on random splits, colored by ARG class with train datapoints opaque and test data points outlined. **c**, t-SNE plot of CLS token embeddings for LR Multiclass model trained on clustered splits, colored by ARG class with train datapoints opaque and test data points outlined. **d**, Bar plot phenotypes recalled from WGS data by tested tools. **e**, Bar plot of ARGs identified by class in WGS data by each tool.

### WGS Results

To demonstrate the usage of *resLen* in a more realistic setting, we used the LR models to analyze a sample WGS data of organisms with laboratory-confirmed resistance phenotypes compiled by VanOeffelen et al.; we applied ResFinder and RGI to the data in a similar manner as points of comparison.^25^ Filtering and mapping classes to those predicted by *resLens* yielded 79 genomes with lab-confirmed resistance phenotypes to between one and three antibiotic classes per organism, resulting in 97 resistance phenotypes to predict across the 83 genomes. Many of the genomes had genes that confer resistance to antibiotics other than those listed in their annotations or selected in our sampling, likely due to the laboratory testing being non-exhaustive. Additionally, these genomes did not contain annotations as to which gene or genes conferred a given resistance phenotype. As the alignment tools provide information about the best-hit genes in their database for their predictions, we validated the *resLens* predictions by comparing the outputs to NCBI’s non-redundant protein sequence database via blastp and examining results with 100% identity and coverage. We considered a prediction correct if the entry’s page or a literature review of that protein indicated that it conferred resistance to the predicted antibiotic class.

Given these constraints, we examined whether a given tool correctly identified at least one gene that corresponded to a genome’s labeled phenotype(s), as well as how many potential ARGs each method identified. Due to the small sample size, annotation patterns, and the amount of manual work required to validate the *resLens* outputs, this should be viewed as an exploration of how the different tools work on real world data, not a definitive benchmarking of relative performance.

*resLens* and RGI more genes that corresponded to a given phenotype than ResFinder, both in terms of the number of genomes with at least one identified gene corresponding to at least one lab-validated resistance phenotype (97.5%, 98.7%, and 87.3% respectively) and the number of overall phenotypes identified with at least one corresponding identified gene (97.9%, 88.7%, and 80.4% respectively) (**Fig. 4d, Supplementary Table 11**). Each method identified different numbers of potential ARGs, with *resLens* labelling 1,046, RGI 2,079, and ResFinder 514 (**Fig. 4e**).

## Discussion

This study demonstrates the ability of genomic language models to provide classifications of ARGs that are as fast, as accurate, and less database-dependent than alignment and other deep learning tools. While *resLens* models outperformed other deep learning tools and performed competitively with the top alignment methods, several interesting patterns emerged in their relative performance.

The language models performed better than other methods on LR data, but slightly worse than some on SR data. Notably, the performance per ARG class of the *resLens* models positively correlates with the number of training examples for that class, indicating that performance could be further improved by training data augmentation (**Fig. 2c,d, Supp. Table 2)**. Further, the tSNE plots of the final layer CLS token embeddings demonstrates that the model has learned internal representations of the genes that enable generalization (**Fig. 4b**).

Additionally, *resLens* models were amongst the fastest methods evaluated, indicating that deep learning methods need not require additional computational costs compared to classical methods (**Fig. 3b**). Importantly, inference time for deep learning models scales with model size, not training data size, meaning that it could be trained on large datasets without incurring an inference time cost, while alignment algorithms necessarily take longer when run on larger databases. Further optimizations, such as streamlined dataloading methods and dynamic tokenizer padding for specific contexts, could improve inference time further.

The experiments involving retraining the model after holding out specific gene families and with train/test splits clustered by sequence similarity indicate that *resLens* models have the capacity to predict the resistance phenotype of genotypes fundamentally different from those it saw in training, albeit with moderately lower performance. The comparison of the models trained on random and clustered splits in particular validates the performance of the models both in- and out-of-sample. The performance of main model trained on random splits demonstrates that DNA language models can identify ARGs similar to those the training database at the same rate as alignment methods; the only moderate decrease in performance of the model trained on clustered splits demonstrates their ability to generalize to sequences relatively dissimilar to that seen in training while alignment methods necessarily fail. The decrease in performance is almost entirely driven by the binary classification models. In particular, the models trained on clustered splits misclassify non-ARGs as ARGs at a much higher rate than the models trained on random splits. This potentially indicates that the lower diversity of ARG examples in the clustered training data decreases the model’s ability to distinguish ARGs from non-ARGs with related functions, such as components other efflux pumps; this is further suggested by the fact that models trained on clustered data also frequently misclassified multidrug ARGs as non-ARGs. The results demonstrate that the model is not just ‘memorizing’ the dataset, but potentially developing a latent understanding of the mechanisms by which a gene can provide resistance to a specific type of antibiotic.

This conclusion is further supported by examining *resLens*’s false positives from the WGS dataset. For instance, it misidentified an ATP-binding cassette domain-containing protein gene from a *Streptococcus pneumoniae* sample as a gene conferring resistance to macrolide-lincosamide-streptogramin antibiotics. While no literature suggests that this gene confers resistance to this class of antibiotics, many of its structures share extensive similarity with those in a protein controlling resistance to tylosin, a macrolide antibiotic, in *Streptomyces fradiae*, suggesting potential common origin and function.^26^ Using tblastn, the gene’s amino acid sequence had only 28.4% identity with the closest nucleotide sequence in our curated ARG dataset.^27^ This lack of sequence similarity validates that *resLens* is relying on other features in its predictions, potentially involving a latent understanding of protein structure and function.

After these evaluations, the exercise of using *resLens*, RGI, and ResFinder to identify ARGs in unannotated WGS data demonstrates the challenges of comparing across tools and selecting a single ‘best’ tool for a given use case. The three tools, run with approaches and parameters fixed as similarly as possible on the same dataset, yielded counts of ARGs identified by a factor of four. We manually validated *resLens* results by researching the flag genes, finding that of the 1,046 genes flagged, 702 (67.1%) were unambiguous true positives, 246 (23.5%) unambiguous false positives, and 97 (9.3%) ambiguous cases or cases where the model identified an ARG but for the wrong class of antibiotic. While we only manually validated the *resLens* results, it is almost certain that RGI and ResFinder’s genes also contained false positives due to many genes falling near the identify and coverage thresholds of 90% and 60% respectively. Additionally, no patterns are immediately clear in the relative number of genes identified per class for each tool, indicating that behavior is likely context and database-dependent. All of these considerations emphasize the importance of improved database curation and the use of these tools for screening and hypothesis generation, not drawing final conclusions.

Taken together, the performance of the *resLens* models validates the potential for DNA language models to improve bioinformaticians ability to identify ARGs both present and poorly represented in reference databases. They can enable researchers to identify and analyze potential new mechanisms of resistance more rapidly and comprehensively, speeding up the loop between in silico and in vitro experimentation. *resLens*’s performance further indicates that DNA language models could improve detection and analysis of novel genotype-to-known phenotype relationships in other domains.

## Methods

### Data

We initially collected ARGs from two sources: version 2.4.0 of the ResFinder database and RefGene sequences labeled as conferring antimicrobial resistance. Full datasets are available for download from our HuggingFace repository (see Data Availability section).

Given that the datasets varied in their level of granularity in assigning ARG classes to genes, we used the lowest level classifications available across all three datasets; however this turned out to be a relatively high level (e.g., ‘cephalexin’, ‘cephalosporins’, and ‘beta-lactam’ were all coalesced into the class beta-lactam, as cephalexin is a type of cephalosporin, which is a subclass of beta-lactam antibiotics). A full class mapping is available in **Supplementary Table 1**.

We then combined these datasets and removed genes that were either perfect duplicates or perfect subsequences of other genes and that conferred resistance to the same class of antibiotic, prioritizing retaining ResFinder then RefGene. We then retained only sequences that had at least 20 instances in the dataset and fed the data through Prodigal to ensure they all constituted only ORFs, yielding 7,606 ARGs over 12 classes.

To acquire negative examples, we queried GenBank for 100,000 random bacterial sequences between 100 and 6,000bp, approximately the same length distribution as our ARGs. We then aligned these genes with our ARG dataset, removed those with greater than 90 sequences identity with any ARG in our dataset, randomly selected a number equal to that of our ARGs, and performed the same open reading frame finding process with Prodigal to construct the non-ARG class in our dataset.

After initial testing, we supplemented the dataset with synonymous ARG sequences from gene families that *resLens* was frequently predicted incorrectly, as well as with non-ARG sequences that *resLens* was frequently classifying as ARGs. A full table of genes by class is available in **Supplementary Table 2**, and distribution of gene lengths is in **Supplementary Figure 1**.

We used this dataset as is to fine-tune the long-read model. For the SR dataset, we split up the genes into 150bp subsequences, with each inheriting the class label of the original gene. We then randomly selected 80% of each of these datasets for training and validation, and the remaining 20% as a test set. For the SR dataset, sequences derived from the same gene were all assigned to the same split in order to prevent data leakage. We used the same training and test splits for the two stages of each model (see *Language Model Training* below), with non-ARG sequences excluded from the dataset for fine-tuning the second stage model trained to distinguish between different ARG classes.

We created our train, test, and validation splits randomly, rather than by assigning entire gene families to specific splits or clustering by sequence similarity and assigning entire clusters to specific splits as is often done in similar settings. We chose to do this because the primary purpose of *resLens* and the other tools against which it is compared is to recall sequences either in or very similar to those in reference databases. Therefore random train and test splits most fairly evaluate the models’ ability to classify genes with a similar distribution to those in the data seen during training, both allowing us to measure the most relevant part of performance and fairly compare it to other methods, as this is also most analogous to how alignment-based tools operate and the other deep learning models referenced here use this approach. We experimented with retraining the models with clustered splits in order to explore their out-of-sample performance; see the *Novel ARG Classification* below for this analysis.

The WGS dataset consisted of 100 randomly selected sequences from the database assembled by VanOeffelen et al., each of which comes from studies that validated one or several resistance phenotypes of an organism in a lab. As the lab testing was non-exhaustive for all ARG classes and not all were selected in our sampling, many genomes contained ARGs to classes other than the labeled ones. Additionally, the exact mechanisms of resistance or genes implicated were not annotated. These constraints are inherent to any large lab-validated database, so we chose to adapt our evaluation methods to them rather than construct synthetic datasets. Filtering out genomes with resistances to antibiotics that *resLens*, RGI, or ResFinder did not predict left 79 genomes with 99 resistance phenotypes. The distribution of resistance phenotypes is shown in **Fig. 1c**. Each file contained several hundred contigs ranging from 1 kbp to 100 kbp.

### Language Model Training

The base model for *resLens* is a DNA Transformer encoder model. For inference, it performs the following: 1) divides a DNA sequence into the subsequences, or tokens, that most efficiently compress the sequence, 2) assigns embeddings to each token, 3) updates the token embeddings via attention modules, 4) passes the final embeddings through a multilayer perceptron classifier to assign a class. Parameters used in steps 1, 2, and 3 are updated during pretraining, while those in stages 2, 3, and 4 are updated during fine-tuning. See **Fig. 1b** for details on the pretraining, fine-tuning, and inference processes.

The specific model chosen was a seqLens 89M parameter model developed by Baghbanzadeh et al. This model has a DeBERTa-v2 base, meaning that it employs disentangled attention, which means that content and position embeddings are treated separately by the attention mechanisms, rather than combined in other commonly used BERT-based models used for biological sequence data, such as Nucleotide Transformer^18^ and ESM-2^28^. Additionally, it uses byte pair encoding for tokenization, which dynamically learns the most common combinations of nucleotides in the pretraining data, enabling more efficient and biologically meaningful tokenization than k-mer tokenization.

We finetuned two models each for the LR and SR versions of *resLens*, one binary classification model to distinguish between ARGs and non-ARGs, and one multiclass classification model to categorize those sequences that the first model had identified as ARGs. Each model was fully fine-tuned for 10 epochs with a learning rate of 1e-5, a batch size of 8, and 2 gradient accumulation steps. We conducted ten-fold cross-validation, recorded model performance on the validation set every 500 steps, and saved a checkpoint of the best model observed by validation weighted F1 during the training process; this best model was used to evaluate performance on the test set. All training was conducted on NVIDIA Tesla V100 GPUs at the George Washington University High Performance Computing cluster.

### Long and Short Read Evaluation

We evaluated the models against the test LR and SR datasets in two different ways. First, we evaluated each *resLens* model on the appropriate version of the test dataset, meaning that we scored the binary model on its ability to distinguish ARGs and non-ARGs in the test dataset and the multiclass ARG model on its ability to distinguish between different types of ARGs in the test dataset, with non-ARGs excluded. Second, we evaluated *resLens* and the selected other tools on a ‘full pass’ of the test dataset, meaning that we had the *resLens* binary model predict which genes were ARGs and then had the multiclass ARG model make predictions for only the genes the binary model had classified as ARGs. We ran all GPU-enabled methods on a single NVIDIA Tesla V100 GPU, and all others were parallelized on Dual 20-Core 3.70GHz Intel Xeon Gold 6148 processors. Times were calculated using wall clock time; they include all data preprocessing steps, but not any database download or retrieval steps. We used the default parameters as of the most recent versions of each tool on 28 January, 2026, with the following exceptions:

- AMR++: We set the -min 100 -max 100 -samples 1 in order to get the results on one pass, the entire dataset rather than bootstrapped subsamples.
- KARGA: s:123 to set the random seed for reproducibility
- Meta-MARC: -l 1 -m to get the level of antibiotic classes that are most closely aligned with ours and to return only one hit per read
- ResFinder: -l 0.1 for the SR data to account for the length of the sequences and -ignore_missing_species for both models due to our lack of consistent species information
- RGI: -include_loose for the loose analysis and -low_quality for the SR analysis.

AMR++ does not provide gene-level annotations but counts of genes associated with specific phenotypes. We compared it to other models by using these counts and the labels from resLens, KARGA, and RGI-Loose to create predicted distributions of the classes in the LR test dataset, then compared it to the true distribution of classes in the test dataset via Jenson-Shannon divergence and Spearman correlation (**Supplementary Table 7**).

### Novel ARG Classification

To evaluate the ability of this methodology to predict the phenotype of novel or unseen ARGs, we performed two analyses. For the first experiment, we re-fine-tuned the foundation model with certain gene families removed from the dataset, then evaluated the ability of the new model to classify and cluster genes from the heldout families. Second, we constructed new train, validation, and test splits by clustered sequences by sequence identity then sorted the clustered into splits, ensuring that no genes from one split share substantial similarity with any in another.

For the former, we searched our long-read dataset for gene families that were 1) in ARG classes that were well-represented in our data and had a high diversity of gene families conferring resistance and 2) had low sequence similarity to other gene families that also conferred resistance to the same class of antibiotics. We relied on the gene family annotations from the ResFinder and RefGene databases for this search. This resulted in us selecting the blaADC gene family of class A beta-lactamases from the ARGs that conferred resistance to beta-lactam antibiotics and the aminoglycoside nucleotidyltransferase (ANT) gene family from the ARGs that conferred resistance to aminoglycoside antibiotics. The blaADC sequences had a maximum of 61.6% and a median of 60.4% identity with the closest other genes conferring beta-lactam resistance; the ANT sequences had a maximum of 64.8% and a median of 50.9% to the closest other genes conferring aminoglycoside resistance. We then removed the genes in these two families from the dataset and placed them in a separate holdout set. Additionally, we removed any genes in the beta-lactam and aminoglycoside ARG classes that lacked a gene family annotation from the remaining dataset in order to eliminate the possibility of data leakage. **Supplementary Table 12** details the breakdown of the remaining datasets and heldout genes.

Removing the blaADC sequences posed an easier version of the problem of classifying novel ARGs, as there were other class A beta-lactamase genes in the training dataset; despite their modest sequence similarity, all class A beta-lactamase genes confer resistance via similar mechanisms of serine hydrolysis. On the other hand, removing the ANT sequences posed a harder version of the problem as they both had lower sequence similarity with and a fundamentally different mechanism of resistance to other aminoglycoside ARGs.

We fine-tuned the same foundation models for multiclass ARG classification with the same hyperparameters and the main models, but this time for only one fold, with 20% of the training data used as a validation set and the best model selected via weighted F1 score on this set. We then evaluated these models’ performance on the two heldout gene families.

For the second evaluation method, we used CD-HIT-EST to create clusters from our original dataset of ARGs and non-ARGs with 90% sequence identity. We then assigned these clustered into train and test splits, using a stratified approach to ensure that the classes were represented in each set in approximately equal proportion. We split the LR sets into SR sets by the method as before. We then finetuned the base model with the same procedure as before to create new binary and multiclass LR and SR models and evaluated them on the test set.

### WGS Evaluation

Due to the Whole Genome Sequencing (WGS) data coming from non-exhaustively tested organisms and lacking annotation of specific genes or mechanisms conferring a given resistance phenotype, we focused our analysis on 1) the recall rate for the lab-validated phenotypes, 2) the amount of the resistome correctly identified, and 3) the number of hits returned, with an in depth manual validation of those returned by the *resLens* models.

We evaluated the *resLens* LR models, fully re-fine-tuned on our entire dataset to ensure that the models had the opportunity to learn all of the resistance gene families, and ResFinder on the 79 genomes we selected. We used Prodigal to identify ORFs in the genomes, then used the two-stage *resLens* models to make predictions on each ORF. We wanted to restrict the *resLens* output to only high-confidence predictions, so we used only those predictions that were in or above the 90% percentile in prediction probability for each ARG class. We ran ResFinder and RGI with their default parameters for assembled contigs, provided the species name when available, and restricted our analysis to those antibiotic classes that *resLens*, ResFinder, and RGI predict for fair comparison.

We validated the *resLens* predictions by comparing the outputs to NCBI’s non-redundant protein sequence database via blastp and examining results with 100% identity and coverage. If the entry’s page or a literature review of that protein indicated that it conferred resistance to the predicted antibiotic class, we labeled it as correct. In some cases, proteins confer resistance to multiple antibiotic classes or have mixed evidence, which we outline in **Supplementary Table 10**. Only those genes with unambiguous evidence or resistance to the correct class were marked as correct predictions for our analysis. We did not analyze RGI and ResFinder’s results for their accuracy due to resource constraints.

## Data Availability

All of the data used for training and testing the models, as IDs to download the whole genome sequencing data from PATRIC, is available at https://huggingface.co/collections/omicseye/reslens-6834aae78d6a59c46d156744.

All of the code used to train the models and perform inference is available at https://github.com/omicsEye/resLens.

## Acknowledgments

This work was supported by the National Science Foundation grant 2109688 to AR and KAC. We are grateful to the GW High Performance Computing team for providing outstanding resources and support for this work.

## Author contributions

A.R. and M.M. conceived the method; M.M. implemented the approach; M.M. tested and packaged the software and evaluated the performance; M.M. and A.R. provided online documents and software. M.M. and A.R. analyzed application datasets. KAC and KD guided data interpretation and result structuring. M.M. and A.R. drafted the manuscript. A.R. and K.C. acquired funding for the project. All authors discussed the results and wrote the manuscript.

